# Theoretical and Empirical Performance of Pseudo-likelihood-based Bayesian Inference of Species Trees under the Multispecies Coalescent

**DOI:** 10.1101/2025.01.28.635282

**Authors:** Nicolae Sapoval, Zejian Liu, Mehrdad Tamiji, Meng Li, Luay Nakhleh

## Abstract

Likelihood-based inference under the multispecies coalescent provides accurate estimates of species trees. However, maximum likelihood and Bayesian inference are both computationally very demanding. Pseudo-likelihood has been previously proposed as a computationally efficient alternative to full phylogenetic likelihood calculations in the context of maximum likelihood estimation. However, theoretical and practical aspects of pseudo-likelihood in the context of Bayesian inference have not been explored. In this work, we provide strong theoretical guarantees and an empirical evaluation of pseudo-likelihood-based Bayesian inference of species trees under the multispecies coalescent. Our contributions are threefold. First, we prove a bound on the convergence rate for species tree topology inference and a Bernstein-von Mises result for branch lengths under model misspecification. Second, we provide an empirical comparison of full- and pseudo-likelihood-based Bayesian inference on synthetic data. Finally, we demonstrate the practical scalability of pseudo-likelihood-based inference by analyzing two biological datasets.

## 1 Introduction

Phylogenomic inference of the evolutionary history of a set of species from genomic data has become commonplace due to its ability to incorporate genome-wide signals that help resolve evolutionary relationships that are not easy to elucidate otherwise [5, 7]. However, this inference is very challenging owing in large part to phylogenetic discordance—the phenomenon where evolutionary histories of different genomic regions disagree with each other as well as with the evolutionary history of the populations from which the genomes are obtained. Biological processes that could cause this phenomenon include incomplete lineage sorting (ILS), gene duplication and loss, and horizontal gene transfer [17, 21].

The multispecies coalescent (MSC) [27, 6] has emerged as a powerful mathematical model for inference of species trees from multi-locus data while accounting for ILS. Furthermore, significant progress has been made in developing species tree inference methods under the MSC [18]. In practice, likelihood-based approaches such as maximum likelihood [14, 38, 25] and Bayesian inference [3, 23] provide high accuracy. Furthermore, Bayesian inference allows for incorporating prior knowledge and quantifying uncertainty in the inference results—two advantages over maximum likelihood method and the (much faster) summary statistic methods [18].

While MSC-based likelihood calculations using bi-allelic marker data can be done efficiently [3], that is not the case when the data consists of gene tree estimates, which has limited the applicability of likelihood-based inferences to datasets whose sizes do not exceed tens of species and hundreds of loci [24]. Thus, significant efforts have been made to accelerate Bayesian inference methods by introducing new sampling proposals [8, 41] or restricting the space of explored trees [34]. In all of these cases, the full likelihood calculations still remain a bottleneck, since every iteration of the Markov chain Monte Carlo (MCMC) sampling algorithm requires evaluation of the full likelihood ratio for the proposal. To address this challenge, pseudo-likelihood was introduced as a computationally efficient alternative to full likelihood computation in the context of maximum likelihood inference [16]. Furthermore, the authors proved that the maximum pseudo-likelihood estimate is statistically consistent under the MSC [16] model.

Given the added advantages of Bayesian inference (which include incorporating prior information, estimating parameters beyond the tree topology, and quantifying uncertainty), it is important to investigate the question of whether the use of pseudo-likelihood instead of full likelihood in a Bayesian setting under the MSC improves scalability while achieving theoretical guarantees on the posterior distribution. To our knowledge, this question has not been explored and this study aims to address it.

The contributions of this work are as follows. We first prove theoretical guarantees for frequentist species tree topology estimation under pseudo-likelihood, which refines the known result of [16] with improved utility for practitioners. Unlike the non-Bayesian focus on point estimation, Bayesian inference requires characterizing the entire posterior distribution, which must account for the model misspecification inherent in pseudo-likelihood methods. Here, we address this challenge from a theoretical perspective to ensure broad applicability. In particular, we show that, under mild conditions, the pseudo-likelihood-based posterior distribution of branch lengths converges to a normal distribution with particular mean and covariance parameters. This characterizes the behavior of pseudo-likelihood-based posterior distributions and is achieved by proving a so-called Bernstein-von Mises (BvM) result under model misspecification. Our BvM result yields insights into the coverage of credible intervals and the consistency of the posterior distribution. In addition to our general theory, we provide empirical comparisons between full- and pseudo-likelihood-based Bayesian inference of species trees. We compare the two approaches on simulated datasets in terms of the accuracy of topology estimation and agreement in the inferred posteriors and species tree distributions. We also explore the computational speedup offered by pseudo-likelihood based inference. Finally, we explore scalability of pseudo-likelihood-based Bayesian analyses to biological data from spiders [7] and turtles [5].

It is important to emphasize here that summary statistic methods, such as ASTRAL [19], are much more scalable, yet provide less information, than Bayesian or maximum likelihood methods. The goal of this work is not to provide a method that outperforms methods like ASTRAL in terms of scalability of species tree topology inference. Our work, instead, is aimed at adding important theoretical and empirical results to the literature on Bayesian inference of species trees under the multispecies coalescent and providing practitioners with a new tool for phylogenomic analyses.

Pseudo-likelihood-based Bayesian inference of species trees is implemented as function `MCMC_GT` with the flag-pseudo in the software package PhyloNet [32, 37].

## 2 Methods and Theoretical Results

### 2.1 Phylogenetic pseudo-likelihood function

For the remainder of the manuscript, we will use *N* to denote the number of taxa and *M* to denote the number of gene trees. Let *S* be a rooted species tree and let 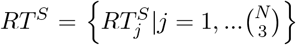 be the set of rooted triples in *S* (i.e., rooted 3-taxon subtrees of *S*; see Fig. 1). Given a rooted triple 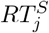, we associate with it a corresponding internal branch length *b*_*j*_, and we will call the set of all such internal branch lengths 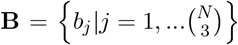 (e.g., in Fig. 1, we have *b*_1_ = *x, b*_2_ = *x* + *y, b*_3_ = *b*_4_ = *y*).

**Figure 1:**
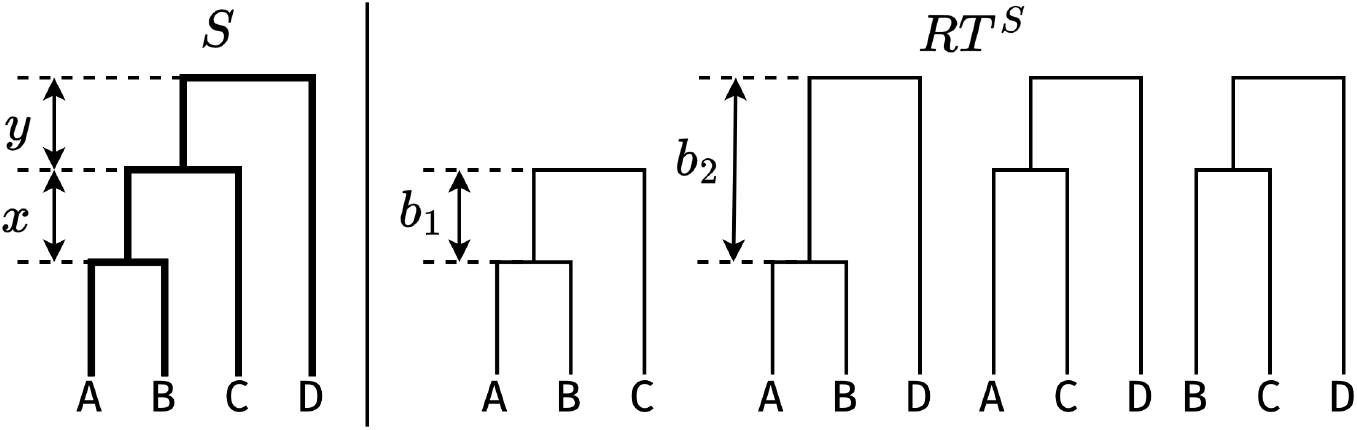
A 4-taxon species tree *S* (left) and corresponding set of rooted triples *RT* ^*S*^ (right). Each rooted triple has an associated internal branch length *b*_*j*_.

If we consider an arbitrary rooted triple 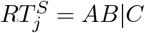 (i.e., *A* and *B* coalesce first; see leftmost 3-taxon tree in Fig. 1), then under the coalescent process the probability that *ab*|*c* occurs in a gene tree generated from *S* is 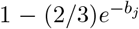, and the probabilities of *ac*|*b* and *bc*|*a* are both equal to 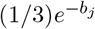. Let *x*_*j*1_, *x*_*j*2_, and *x*_*j*3_ denote the counts of triples *ab*|*c, ac*|*b*, and *bc*|*a* observed in the gene trees (note that we have *x*_*j*1_ + *x*_*j*2_ + *x*_*j*3_ = *M*). Then it follows that the counts follow a multinomial distribution

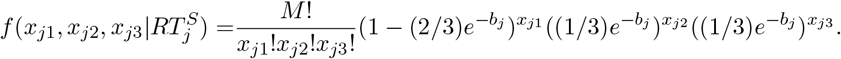

Now, we can define the pseudo-likelihood of a species tree *S* given a collection of gene trees *G* as

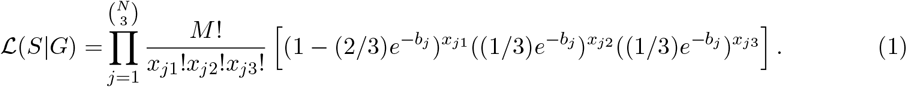

For example, consider the species tree *S* on four taxa and let *G* = {*G*_1_, *G*_2_, *G*_3_} be three gene tree topologies arising from *S* (Fig. 2). Then the full log-likelihood (*ℓ*_*full*_) for observing *G* given *S* with branch lengths *x* = 0.5 and *y* = 0.6 is

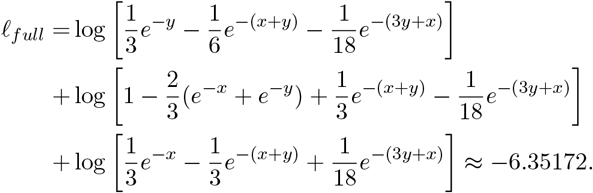

**Figure 2:**
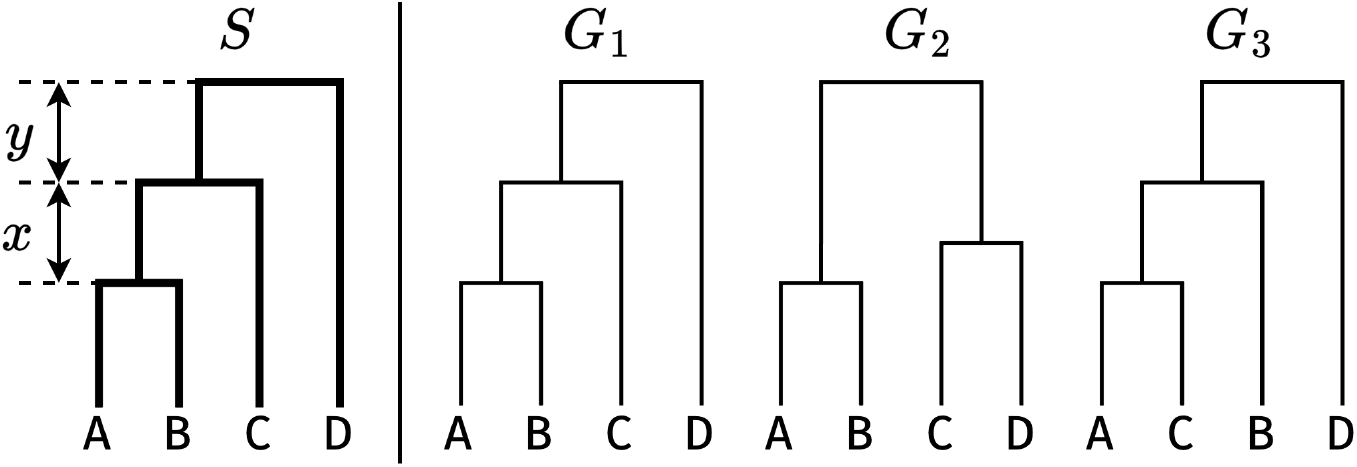
A 4-taxon species tree *S* (left) and corresponding collection of gene tree topologies *G* = {*G*_1_, *G*_2_, *G*_3_} (right).

In contrast, the log-pseudo-likelihood (*ℓ*_*pseudo*_) in this case is equal to

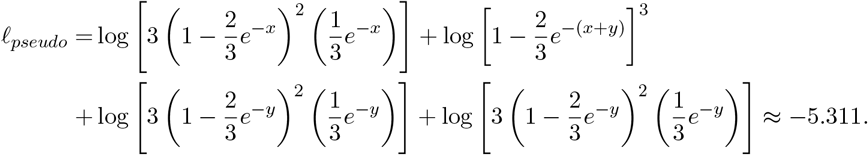

### 2.2 Pseudo-likelihood-based Bayesian inference

For the Bayesian inference under pseudo-likelihood we employ a standard Metropolis-Hastings algorithm where in the acceptance ratio calculation we substitute the value of the likelihood ratio of the proposal and current state by the corresponding pseudo-likelihood ratio. Note that since for a fixed collection of gene trees the value of 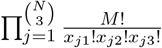 in Eq. (1) is constant, it follows that the ratio can be simplified to

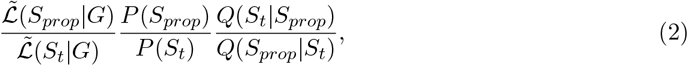

where

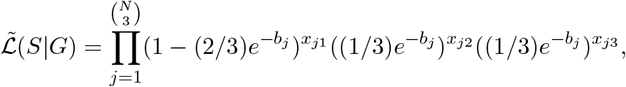

*P* (*S*) is the prior, and *Q*(*S*|*S*_*t*_) is the proposal probability. For the numerical stability of the calculation logarithms of all the ratios are used in the PhyloNet [32, 36] implementation.

### 2.3 Theoretical results

Let the topology of a true species tree be *S*_0_ and the corresponding set of branch lengths **B**_0_. We use Ŝ to denote the estimated species tree topology by maximizing the pseudo-likelihood function. For any set of branch lengths **B**, we denote the minimum enclosed branch length as min **B**. Throughout this paper, the Euclidean norm is denoted by ∥ · ∥.

We first refine the result of Liu *et al*. [16] by defining an explicit convergence rate for the topology estimation in terms of *N*, *M*, and a branch length parameter of the true species tree. This result holds in finite-sample settings, i.e., for any finite number of taxa *N* and number of gene trees *M*.

#### Theorem 1

(Convergence rate of topology estimation). *The probability that the tree topology Ŝ estimated from a set of gene trees G under the MSC, matches the true species tree topology S*_0_ *is bounded by*

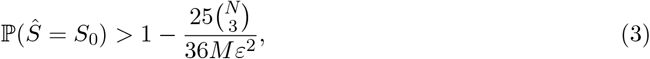

*where* 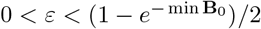.

*Proof of Theorem 1*. In this proof, we use *n* to denote the sample size, i.e., the number of gene trees *M*. Let *B*_0,*j*_ denote the branch length corresponding to the taxa triple *j* in the true species tree *S*_0_. We define a set of events as follows:

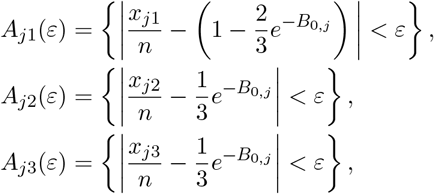

for 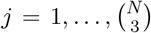, and we use the notation *A*^*c*^ to denote the complement of the set *A*. We can then establish a bound on the probability that the maximum pseudo-likelihood correctly estimates the true topology:

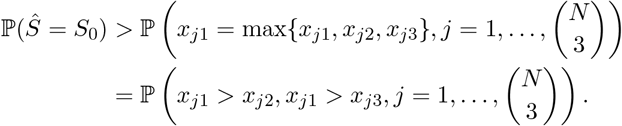

Under events *A*_*j*1_(*ε*), *A*_*j*2_(*ε*) and *A*_*j*3_(*ε*), it follows that *x*_*j*1_ *> x*_*j*2_ and *x*_*j*1_ *> x*_*j*3_ if

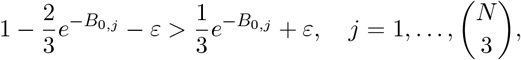

i.e., *ε* is any number satisfying 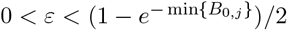. Hence,

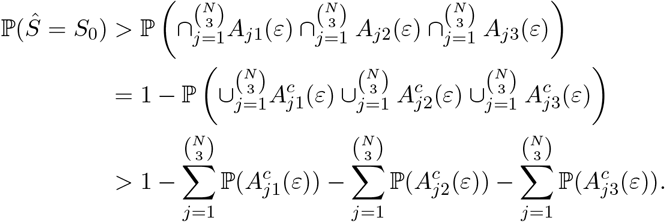

By Chebyshev’s inequality,

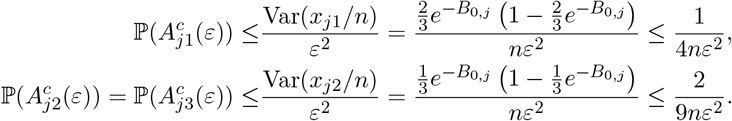

Therefore,

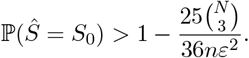

#### Remark 1.

*Convergence rate of Ŝ has been studied in [16]*. *We obtain a refined error bound by revising the proof in* (3) *and additionally specify the required range of ϵ, as opposed to “a small positive real number” described in [16]*. *This explicit range in* (3) *leads to practical guidance on its optimal value for accurate topology estimation. For example, setting ϵ to its upper bound maximizes the error bound for the probability of Ŝ matching the true topology S*_0_. *In particular, if we let p denote a target probability of this event that could be specified by users, then the minimum number of gene trees needed to achieve this probability is given by* 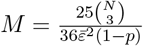, *where* 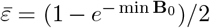 *Also, for any N, letting M* → ∞ *in* (3) *trivially leads to statistical consistency of Ŝ, i.e., ℙ(Ŝ = S*_0_*) converges to 1*.

We demonstrate the application of the formula in Remark 1 through two examples. First, we fix the number of taxa *N* and illustrate the relationship between the probability of correct identification *p* and the necessary number of gene trees *M*. We select four different numbers of taxa, *N* = 5, 10, 20, 50, and set 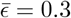. The resulting curves of *p* against *M* for each *N* are depicted in Fig. 3.

**Figure 3:**
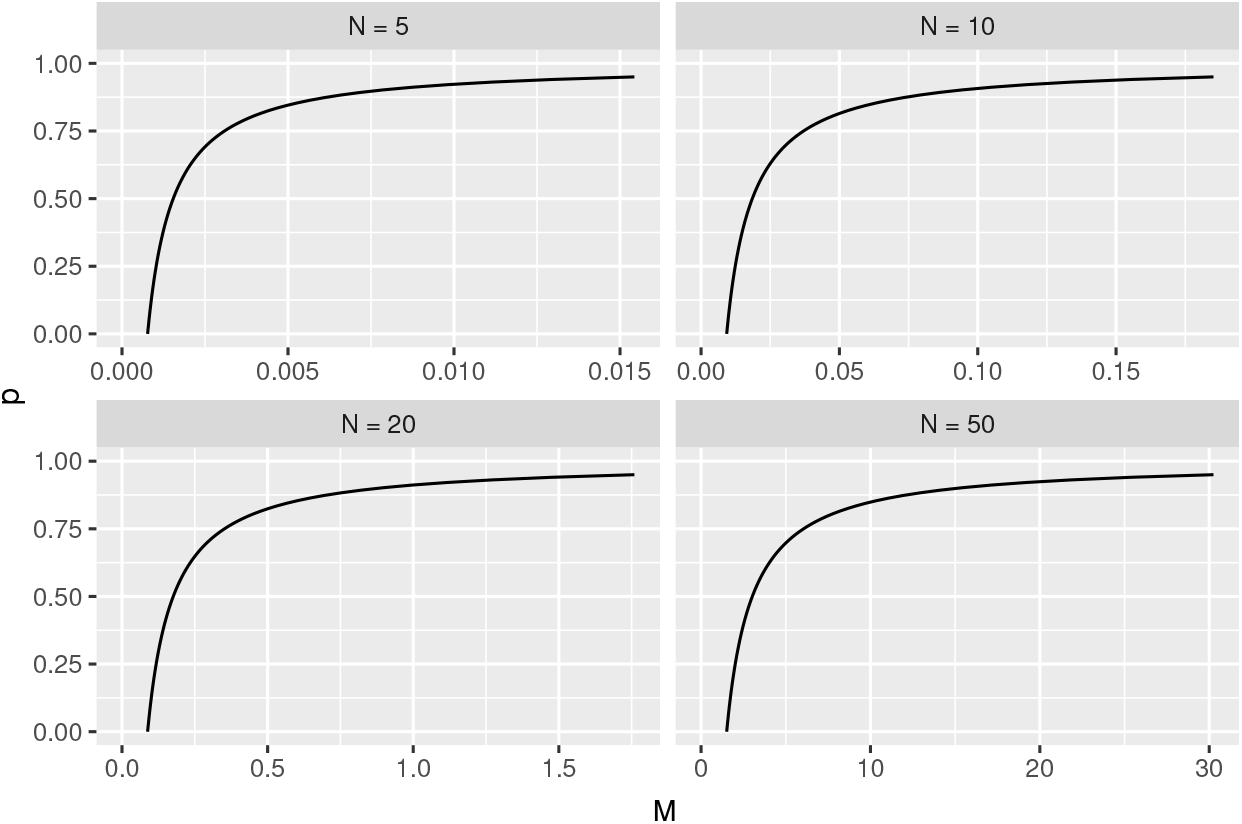
Probability of correct identification *p* against the number of gene trees *M* (×10^5^) for different numbers of taxa *N*.

We then fix the target probability *p* and visualize the relationship between the number of taxa *N* and the number of gene trees *M*. We choose four different target probabilities *p* = 0.5, 0.8, 0.9, 0.95, and fix the value of 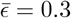. The curves of *N* against *M* for each *p* are depicted in Figure 4.

**Figure 4:**
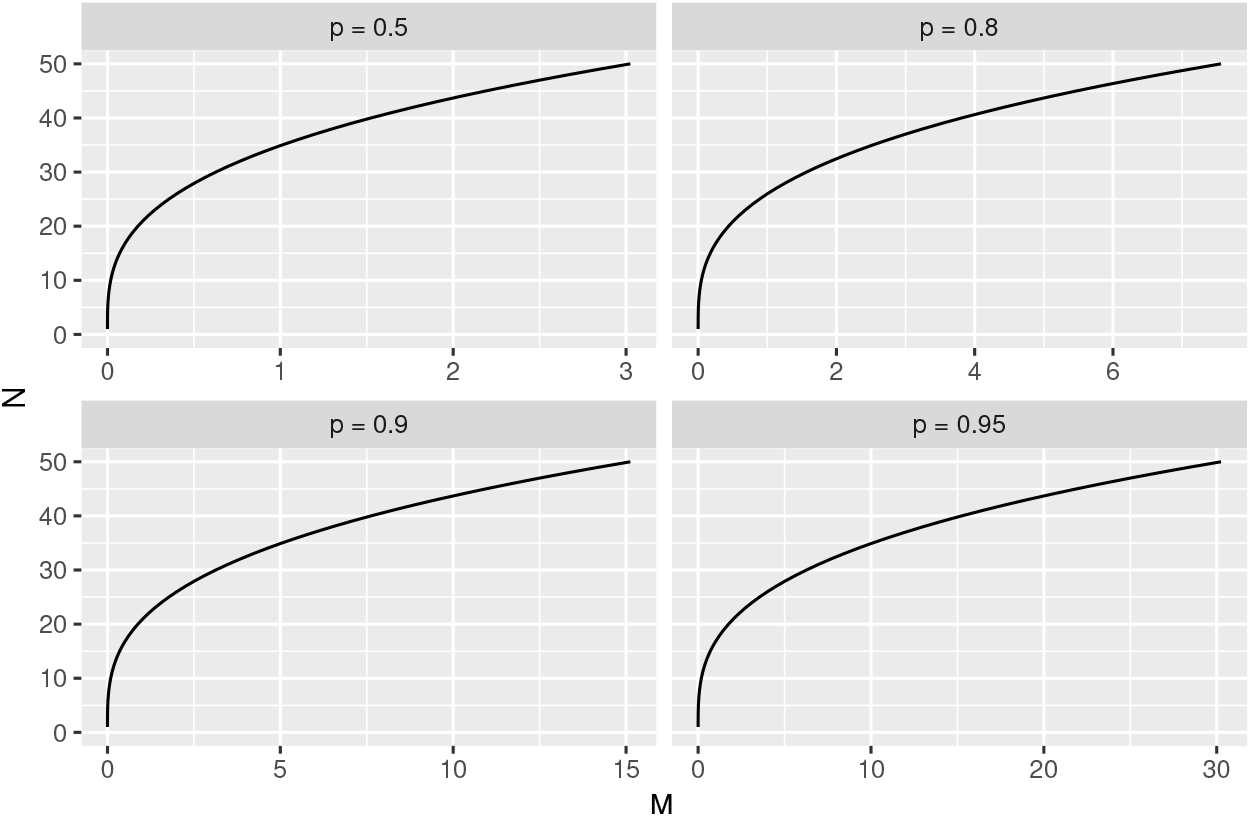
The number of taxa *N* against the number of gene trees *M* (×10^5^) for different probabilities of correct identification *p*.

The finite-sample result in Theorem 1 concerns point estimation. Bayesian implementation based on the pseudo-likelihood would require the prior specification and posterior computation.

Our next theorem studies the behavior of the pseudo-likelihood-based posterior distribution of branch lengths. Since the species tree topology can be estimated accurately with high probability, as established in Theorem 1, and our numerical experiments in Section 3 confirm the accuracy of this estimation using Bayesian methods, we fix the topology at *S*_0_. This choice also simplifies interpretation, as branch lengths are closely linked to the topology. Section 3 also presents empirical results that account for the uncertainty in inferring *S*_0_.

Let Π_*M*_ (**B**|*G*) denote the posterior probability distribution of branch lengths given a collection of gene trees *G*. Our next main result shows that it converges to a normal distribution in total variation distance. Specifically, let ℙ_0_ denote the data generating measure, *p*_0_ denote its associated density relative to the Lebesgue measure, and the density according to the pseudo-likelihood as 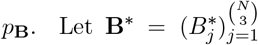 be the branch lengths that minimize the Kullback-Leibler divergence *D*_KL_(*p*_**B**_ ∥ *p*_0_), assumed to exist and be unique, which is a key quantity to describe the behavior of Π_*M*_ (**B**|*G*) due to model misspecification. Let *π*(**B**) denote the prior distribution assigned to **B** with respect to the Lebesgue measure. We assume the following general conditions on the prior, covering a broad class of priors:

(A) The prior density *π*(**B**) has compact support and is continuous and positive in a neighborhood of **B**^*^. Additionally, there exists a constant *c* such that ∥*H*(**B**)∥_2_ ≤ *ce*^∥**B**∥2^, where *H*(**B**) is the Hessian matrix of log *π*(**B**) (i.e., its second-order derivatives) and ∥*H*(**B**)∥_2_ is the 2-norm (spectral norm) of *H*(**B**).

#### Remark 2.

*Assumption A is known as the “prior mass” condition in the statistical Bayesian literature, and is very mild. Note that the support of a distribution refers to the set of all possible values where the probability density is non-zero. Many prior distributions exist to ensure that the density functions are continuous and positive at any point within its support, including* **B**^*^. *The “compact support” condition means that the support of the prior is finite; for each dimension of the prior, one can choose a large enough but still finite interval to truncate a prior distribution supported on the real line. Additionally, Assumption A imposes a limit on the log prior density’s second derivative, a restriction easily met by many distributions that are not heavily tailed. In particular, these assumptions hold for the exponential distribution used for branch lengths in PhyloNet [36] for Bayesian inference of species trees*.

#### Theorem 2

(Bernstein-von Mises under model misspecification for branch length estimation). *Suppose Assumption A holds. Then*,

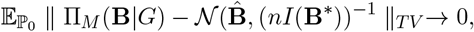

*where*

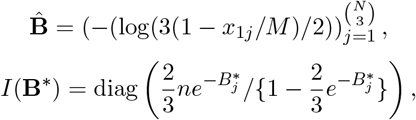

*and* ∥ · ∥_*T V*_ *is the total variation distance used for comparing two probability distributions*.

*Proof of Theorem 2*. In this proof, we use *n* to denote the sample size, i.e., the number of gene trees *M*. We proceed by verifying the conditions established in the literature for Bernstein-von Mises theorems under model misspecification [13, 2], as applied to phylogenetic pseudo-likelihood-based inference:

1. (Prior mass). This is given by Assumption A.
2. (Local asymptotic normality). For every compact set 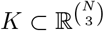, there exist random vectors Δ_*n*,**B**\*_ and nonsingular matrices *V*_**B**\*_ such that

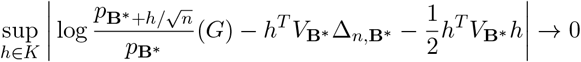

in ℙ_0_-probability.
3. (Consistent testability). For every *ε >* 0 there exists a sequence of tests {*ϕ*_*n*_} such that

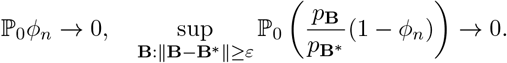

Condition 2 indicates that the frequentist model approaches a normal distribution model asymptotically upon rescaling. We next verify this condition. We first show the consistency of the maximum likelihood estimator (MLE) of the branch lengths under the pseudo-likelihood. Given that the true species tree topology *S*_0_, the log-pseudo-likelihood is

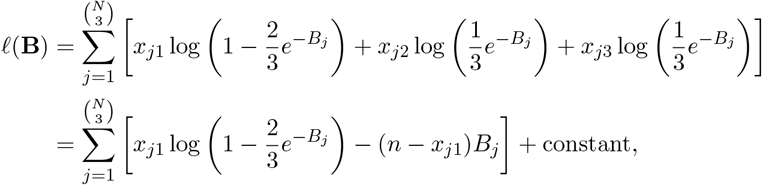

where *x*_*j*1_ = max{*x*_*j*1_, *x*_*j*2_, *x*_*j*3_} and *n* = *x*_*j*1_ + *x*_*j*2_ + *x*_*j*3_. This leads to the MLE for *B*_*j*_ being

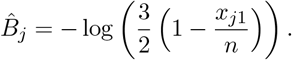

Note that

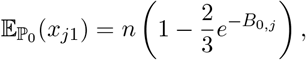

where *B*_0,*j*_ is the length of the internal branch of triple *j* in the true species tree. It can be deduced from the strong law of large numbers that, as the number of genes *M* approaches infinity, the MLE of the branch length converges almost surely to the true value:

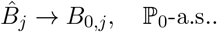

The MLE for **B** based on the pseudo-likelihood is then 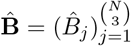, which is a consistent estimator for **B**_0_ ℙ_0_-almost surely. Next we compute the expected value of the score, denoted as Δ_*n*,**B**_:

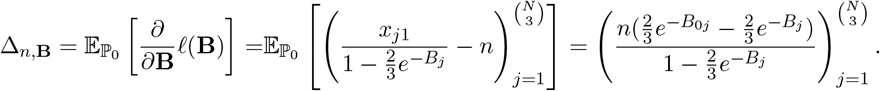

The Fisher information matrix *I*(**B**) is a diagonal matrix, where each diagonal element is

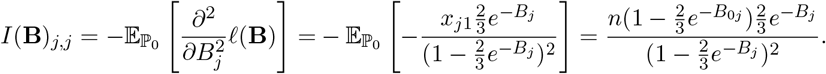

Applying a second-order Taylor expansion to *p*_**B**_, we can validate Condition 2 with Δ_*n*,**B**\*_ = Δ_*n*,**B**_|_**B**=**B**\*_ and *V*_**B**\*_ = *I*(**B**^*^).

Condition 3 posits the existence of consistent tests. It is required that there exists a sequence of uniformly consistent (under ℙ_0_) tests for testing *H*_0_ : **B** = **B**^*^ against *H*_1_ : ∥**B** − **B**^*^∥ ≥ *ε* for every *ε >* 0 based on the pseudo-likelihood model. This condition is mild, and a sufficient condition is that **B** has compact support, which is ensured by Assumption A.

Finally, by Theorem 2.1, Lemma 2.2 and Theorem 3.1 in [13], it holds that

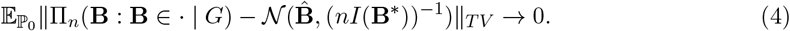

A few remarks are in order.

#### Remark 3.

*Theorem 2 is referred to as a Bernstein-von Mises (BvM) result in the theoretical Bayesian literature. When there is no model misspecification, classical BvM asserts that, under certain conditions, the posterior distribution of a parameter becomes approximately normal and centered at the maximum likelihood estimator as the sample size grows; see, for example, Chapter 10*.*2 of [33]*. *However, pseudo-likelihood-based posterior distribution inherently misspecifies the model by replacing the full likelihood with the pseudo-likelihood. As such, we utilize techniques established in [13] to prove the BvM theorem under model misspecification*.

#### Remark 4.

*The classical BvM phenomenon under the correct model reassures that Bayesian credible intervals are also frequentist confidence intervals. However, this does not hold for the pseudo-likelihood-based posterior distribution due to model misspecification. Let* 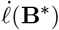 *denote the log-pseudo-likelihood function evaluated at* **B**^*^. *Then the maximum likelihood estimator* 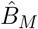 *for the misspecified model has the property that the sequence* 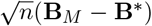 *converges to a Gaussian distribution with mean zero and variance given by* 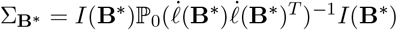.*Covariance matrix I*(**B**^*^) *appearing in the Bernstein-von Mises theorem is not equal to the asymptotic covariance matrix* Σ_**B**\*_, *implying that Bayesian credible sets might not align with confidence sets at the nominal level. Such deviation from classical BvM has been observed and discussed in approximate Bayesian computation contexts, such as variational Bayes [35]*, *and semiparameteric settings with high-dimensional nuisance parameters [15]*. *Our numerical experiments in Section 3 also confirm this in finite samples*.

An implication of Theorem 2 is that the posterior distribution is consistent at **B**^*^, instead of **B**_0_.

#### Corollary 1

(Consistency of posterior). *Under the conditions of Theorem 2*, *then for any ϵ >* 0,

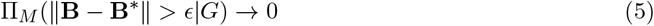

*in* ℙ_0_*-probability*.

*Proof of Corollary 1*. The proof follows directly from the existence of consistent testability demonstrated in the proof of Theorem 2, based on the Schwartz consistency theorem under model mis-specification (Corollary 4.2 in [13]).

## 3 Empirical Results

### 3.1 Simulation and evaluation setup

We simulated species trees using the `SiPhyNetwork` package [11] under the birth-death model with birth-rate *λ* sampled uniformly in the [0.2, 0.5] interval, death-rate *µ* sampled uniformly in the [0.1, *λ*] interval, and hybridization rate set to 0. We simulated a total of 10 species trees for *N* equal to 5 and 10 taxa. For each species tree, gene trees were simulated using `ms` [9] via the `SimGTinNetwork` command in `PhyloNet` [32, 37]. For each species tree, we simulated *M* ∈ {100, 500, 1000, 2500, 5000, 10000} gene trees. For each gene tree, gene DNA sequences of length 1,000 were simulated using Seq-Gen [26] under the Jukes-Cantor model [10] and branch length scaling of 0.01.

We sampled species trees from the posterior distribution using `PhyloNet MCMC_GT` command [36] with total chain length of 1,000,000 steps, burn-in length of 100,000 steps, and sampling frequency of one sample per 1,000 steps. Sampling from the posterior distribution under pseudo-likelihood is now implemented in PhyloNet as part of the `MCMC_GT` command using an additional `-pseudo` flag. For inference from sequence data, gene trees were first inferred for individual genes using IQ-TREE [22] with ModelFinder Plus [12]. The average normalized Robinson-Foulds [28] distance for inferred trees was 0.025 ± 0.077 for 5-species trees and 0.044 ± 0.07 for 10-species trees. Then, species tree inference was performed analogously to the simulated ground truth gene trees scenario described above.

Given that the theoretical analysis indicates that the Bayesian credible sets might not align with the confidence sets at the nominal levels, we assessed the frequentist coverage of the 95% credible sets following the standard methodology [4]. Hence, for each species tree *S*, we generated 100 independent draws of *M* ∈ {100, 500, 1000, 2500, 5000, 10000} gene trees. Then, the pseudo-likelihood MCMC simulation was performed on each of the sets of gene trees yielding 900 samples *Ŝ*_*i*_ from the posterior distribution. Resulting samples were summarized via the `summarize_splits_on_tree` function in `dendropy` (v5.0.1) [20]. For each branch in *S*, we checked if the branch length *θ* fell within the 95% highest posterior density (HPD) credibility interval of the sampled trees Ŝ_*i*_. Finally, we computed the coverage of the 95% credibility interval as the ratio of the branch length values within the intervals to the total number of branch lengths.

In order to assess possible discrepancies between the posterior distributions inferred by pseudo- and full-likelihood-based MCMC samplers, we conducted an additional set of runs for *N* = 5 and the set of simulated gene trees of total length 10,100,000 steps with 100,000 steps used for burn-in. For each of the species tree candidates sampled from the posterior, we recorded its posterior probability and weighted Robinson-Foulds distance [29] to the ground-truth species tree.

## 3.2 Results

### Species tree topology estimation

First, we evaluated pseudo- and full-likelihood-based inference on a set of simulated ground-truth gene trees. The percentage of MCMC samples that had topology matching that of the ground-truth species tree increased as more gene trees were considered (Fig. 5). The pseudo-likelihood-based runs had slightly worse performance for 100 and 500 gene tree settings, but matched the full likelihood results for 5,000 and more gene trees (Fig. 5).

**Figure 5:**
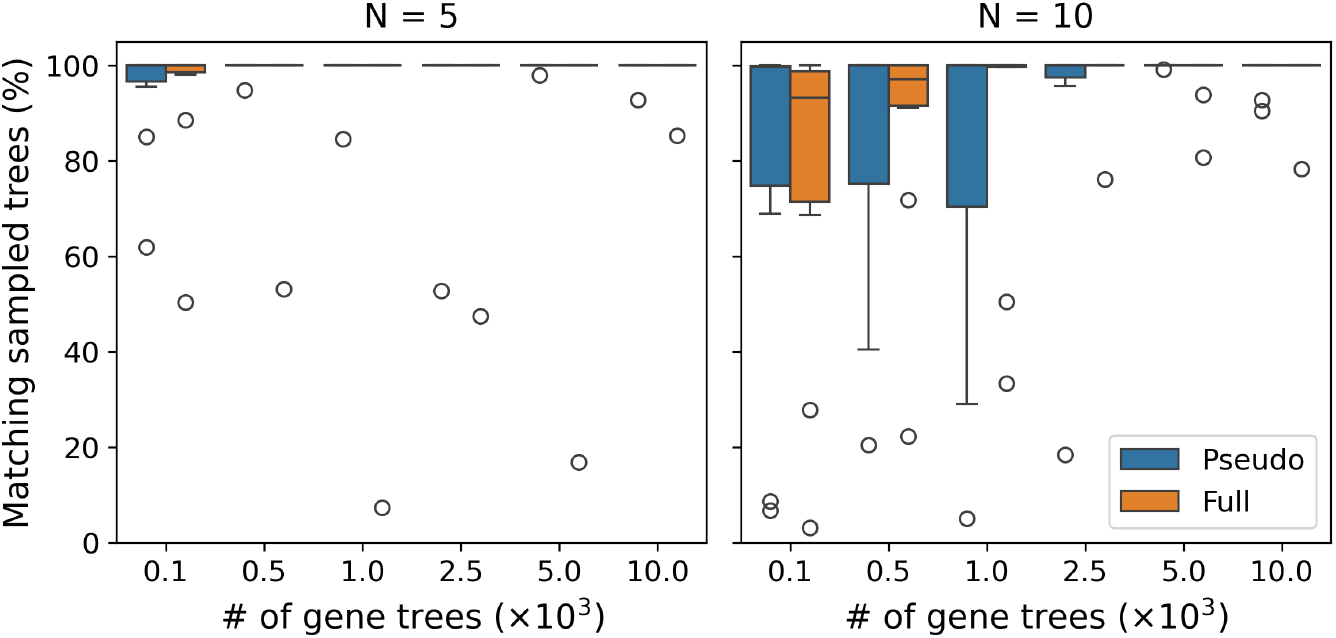
Percentage of the species trees in the pseudo- and full-likelihood-based posterior distribution samples that have correct topology (y-axis) as a function of the number of gene trees (x-axis) and number of species (*N*). The input data consists of the true gene trees.

Next, we evaluated performance of both methods in the presence of potential gene tree estimation errors. After simulating sequences for each gene, the gene trees were inferred and the methods were evaluated with the inferred gene trees as the input. Similarly to the simulated gene trees scenario, as the number of gene trees increased the percentage of sampled trees with correct topology also increased (Fig. 6). Analogously, pseudo-likelihood-based inference had higher variance in the percentage of trees with matching topology for 100 gene trees input (Fig. 6). Both pseudo- and full-likelihood-based inference had a drop in performance when run on inputs consisting of 100 gene trees.

**Figure 6:**
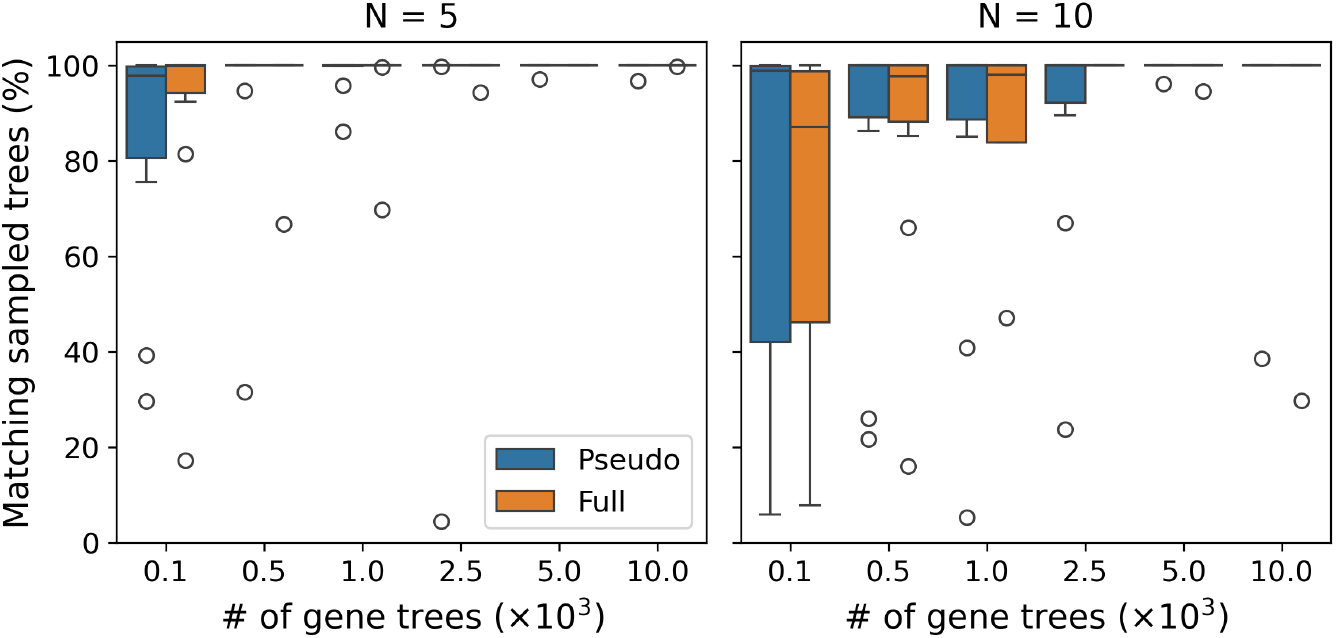
Percentage of the species trees in the pseudo- and full-likelihood-based posterior distribution samples that have correct topology (y-axis) as a function of the number of gene trees (x-axis) and number of species (*N*). The input data consists of the estimated gene trees.

### Branch length credible intervals

Remark 4 implies that the frequentist coverage of the Bayesian credible intervals for the branch length estimates can deviate from its nominal value. We observe that in the empirical setting, the coverage of the 95% Bayesian credibility sets is lower than 95% with median coverage ranging from 0.762 to 0.818 for the 5-species case (Fig. 7, left) and from 0.654 to 0.668 for the 10-species case (Fig. 7, right).

**Figure 7:**
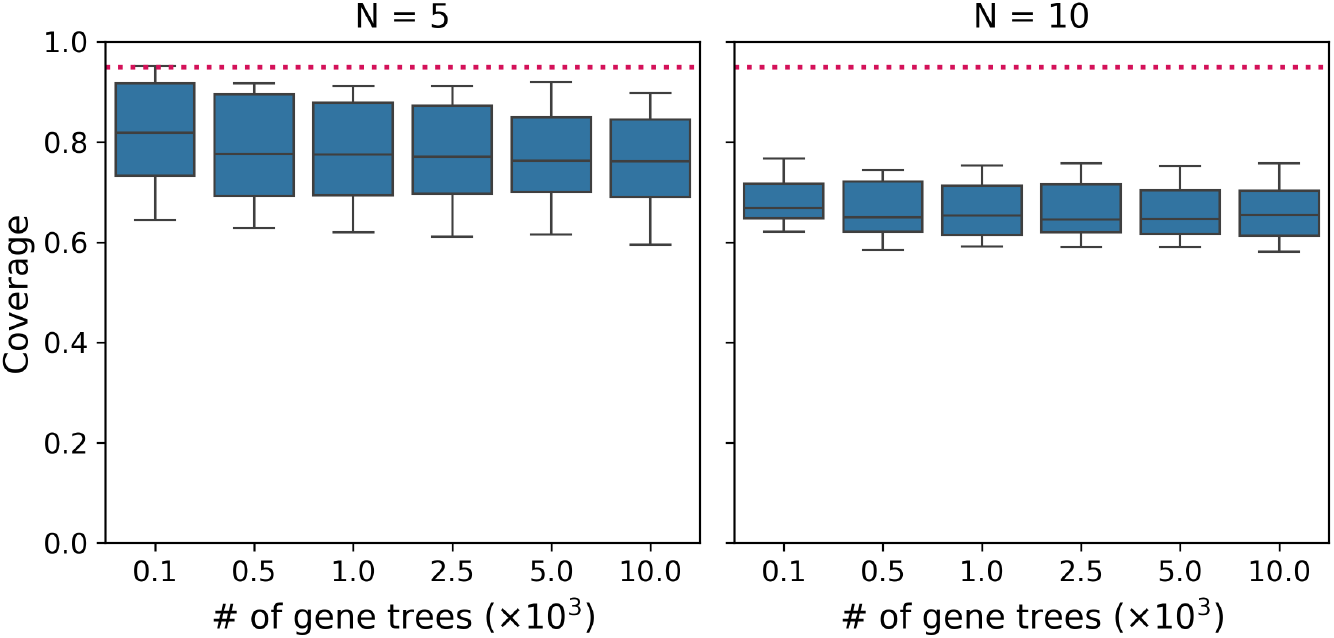
Frequentist coverage of the 95% high posterior probability density intervals for the branch lengths of the species tree. The red dotted line indicates the nominal coverage. Box plots indicate coverage values across 10 replicate species trees. The input data consists of the true gene trees.

We also note that in the 10-species case, there is a reduced variance in the coverage of the credibility intervals.

### Posterior density estimation

To further compare the behavior of pseudo- and full-likelihood-based Bayesian analysis, we compared the distributions of the log posteriors (Fig. 8). Since, the exact values of the log posteriors can differ between the pseudo- and full-likelihood-based analyses by more than an order of magnitude, we centered the distributions by subtracting their mean values.

**Figure 8:**
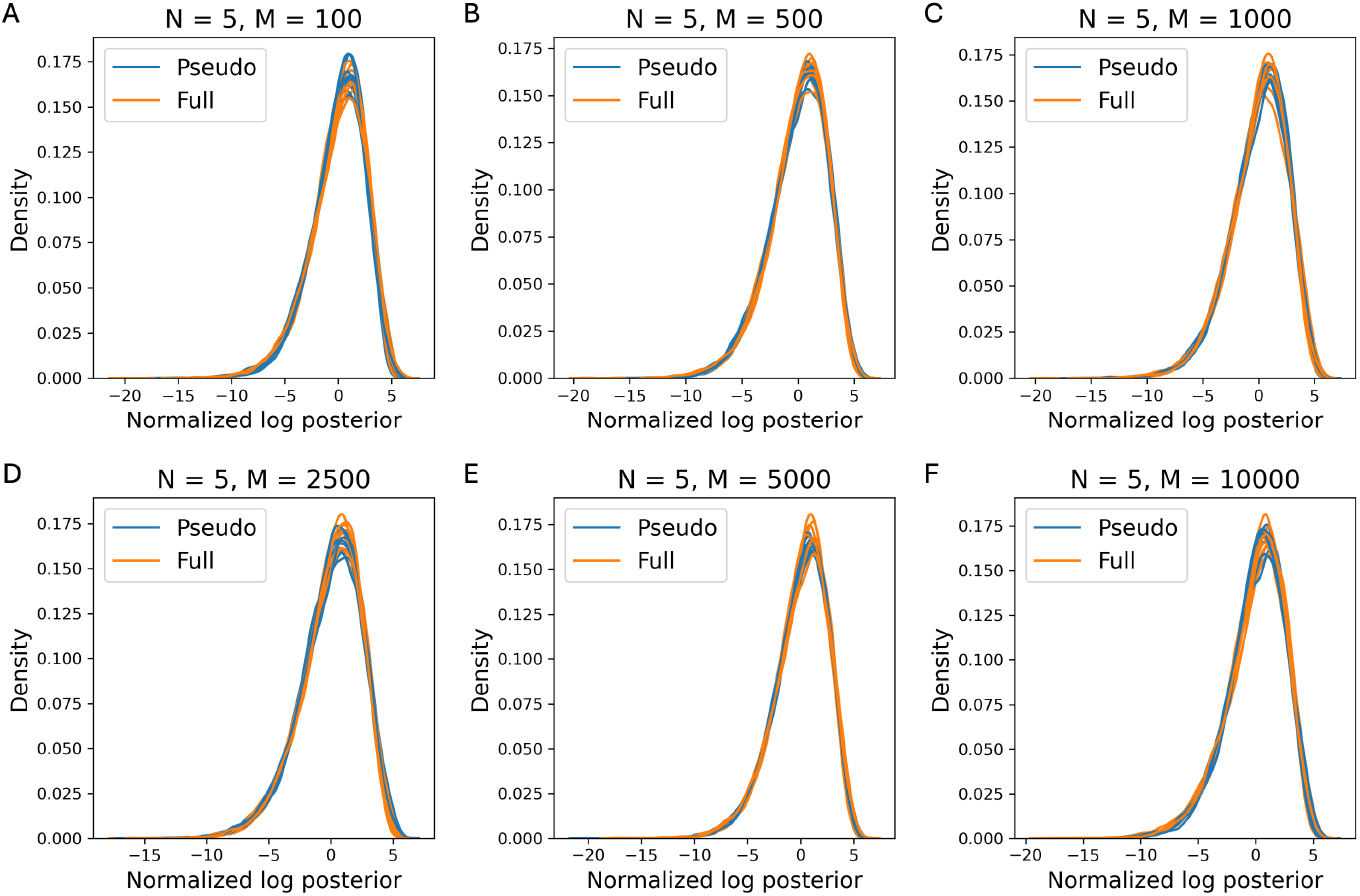
Distribution of centered log posterior probabilities for pseudo- and full-likelihood-based MCMC runs. Each curve represents a single replicate species tree. The input data consists of the true gene trees.

We note that the centered distributions of log posteriors are similar for pseudo- and full-likelihood-based runs, with the *L*_1_ norm of the histogram differences between the distributions ranging from 0.026 to 0.130 with median value of 0.061. We also compared distributions of weighted Robinson-Foulds distances for the inferred trees under pseudo- and full-likelihood-based inference (Fig. 9). Similarly to the distributions of the log posteriors these agree between the pseudo- and full-likelihood-based analyses.

**Figure 9:**
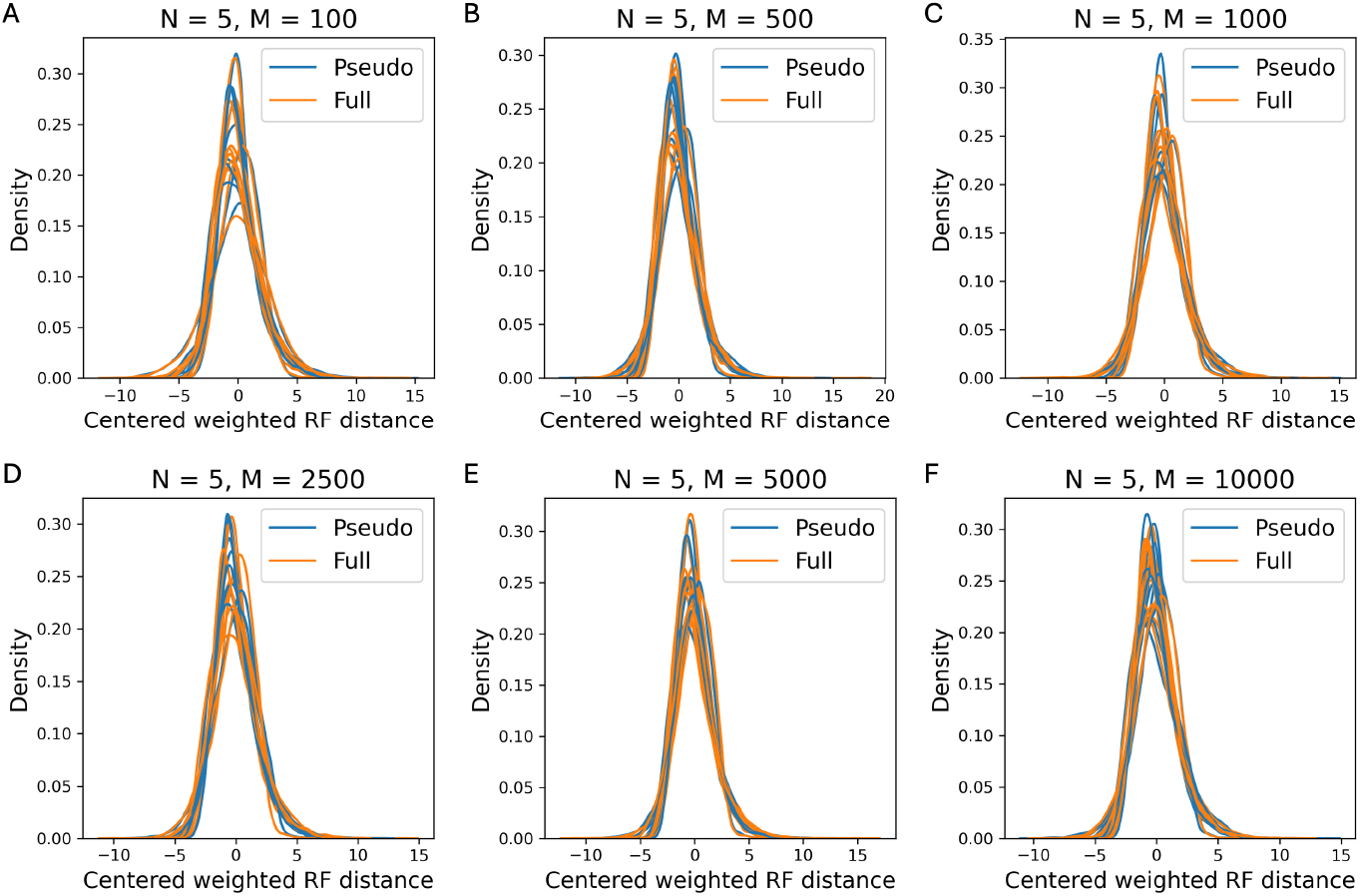
Distribution of centered weighted Robinson-Foulds distances between inferred and true species tree for pseudo- and full-likelihood MCMC runs. Each curve represents a single replicate species tree. The input data consists of the true gene trees.

Furthermore, we note that the acceptance rates of the pseudo- and full-likelihood-based MCMC runs are similar and confined to the [0.4, 0.6] interval (Fig. 10, 11). As the number of gene trees increases, the acceptance rate decreases (Fig. 10, 11), matching the observation that as the number of gene trees increases the proportion of sampled topologies matching the true topology rapidly approaches 1 (Fig. 5, 6).

**Figure 10:**
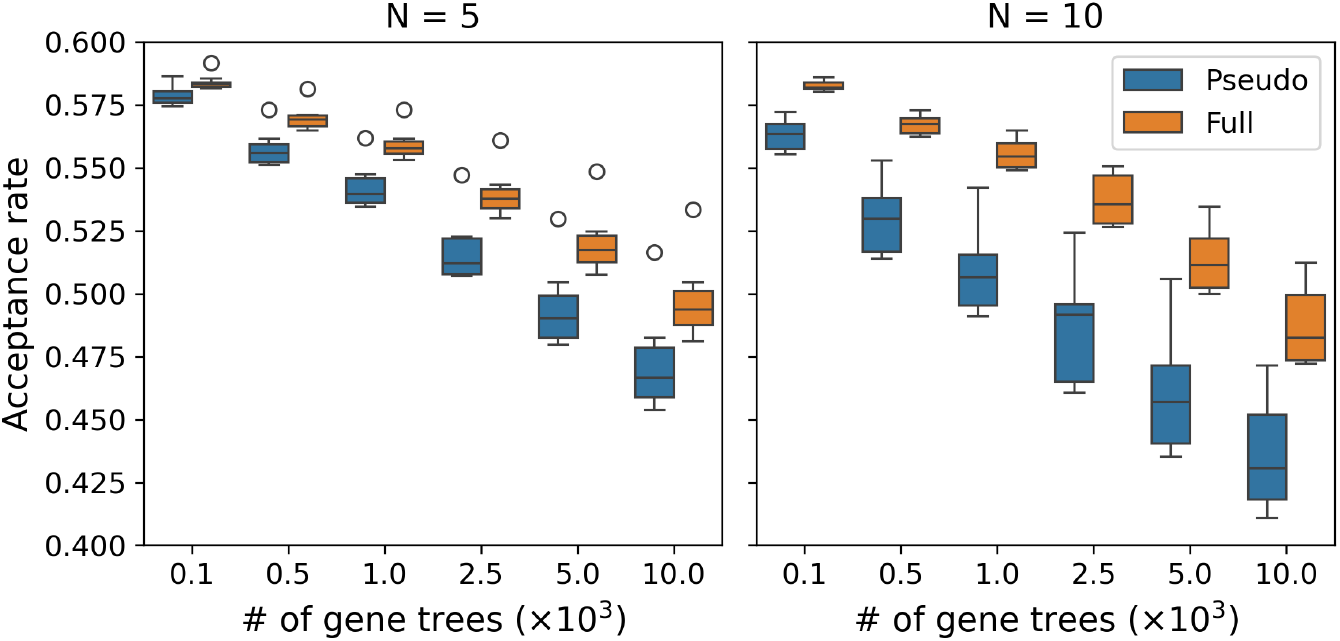
Acceptance rate for the pseudo- and full-likelihood MCMC runs as a function of the number of gene trees (x-axis) and number of species (*N*). The input data consists of the true gene trees.

**Figure 11:**
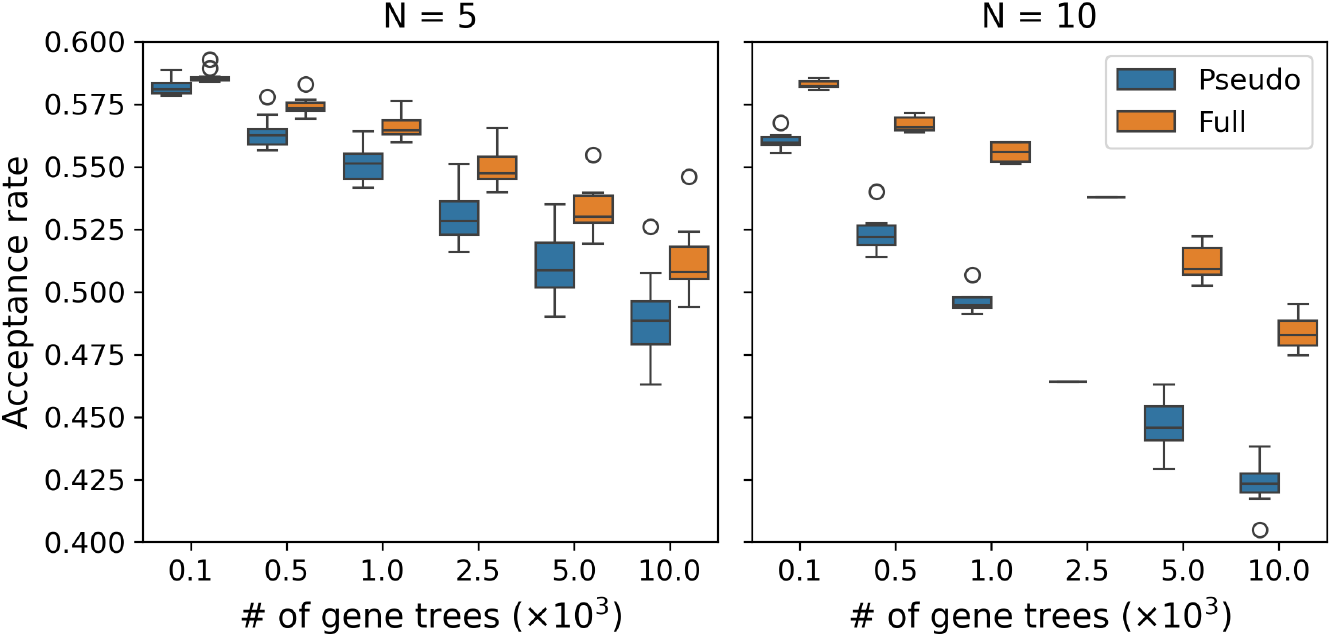
Acceptance rate for the pseudo- and full likelihood MCMC runs as a function of the number of gene trees (x-axis) and number of species (*N*). The input data consists of the estimated gene trees.

### Computational performance

We compared the CPU time required to complete 1,000,000 steps of the MCMC simulation under the pseudo- and full-likelihood functions. For both simulated and inferred gene trees, pseudo-likelihood-based MCMC outperforms full-likelihood-based computation by one to two orders of magnitude (Fig. 12A, B; Fig. 13). In particular, on the datasets with 10-species and *>* 1000 gene trees, pseudo-likelihood-based computation achieves more than 100x speedup compared to the full-likelihood-based analysis.

**Figure 12:**
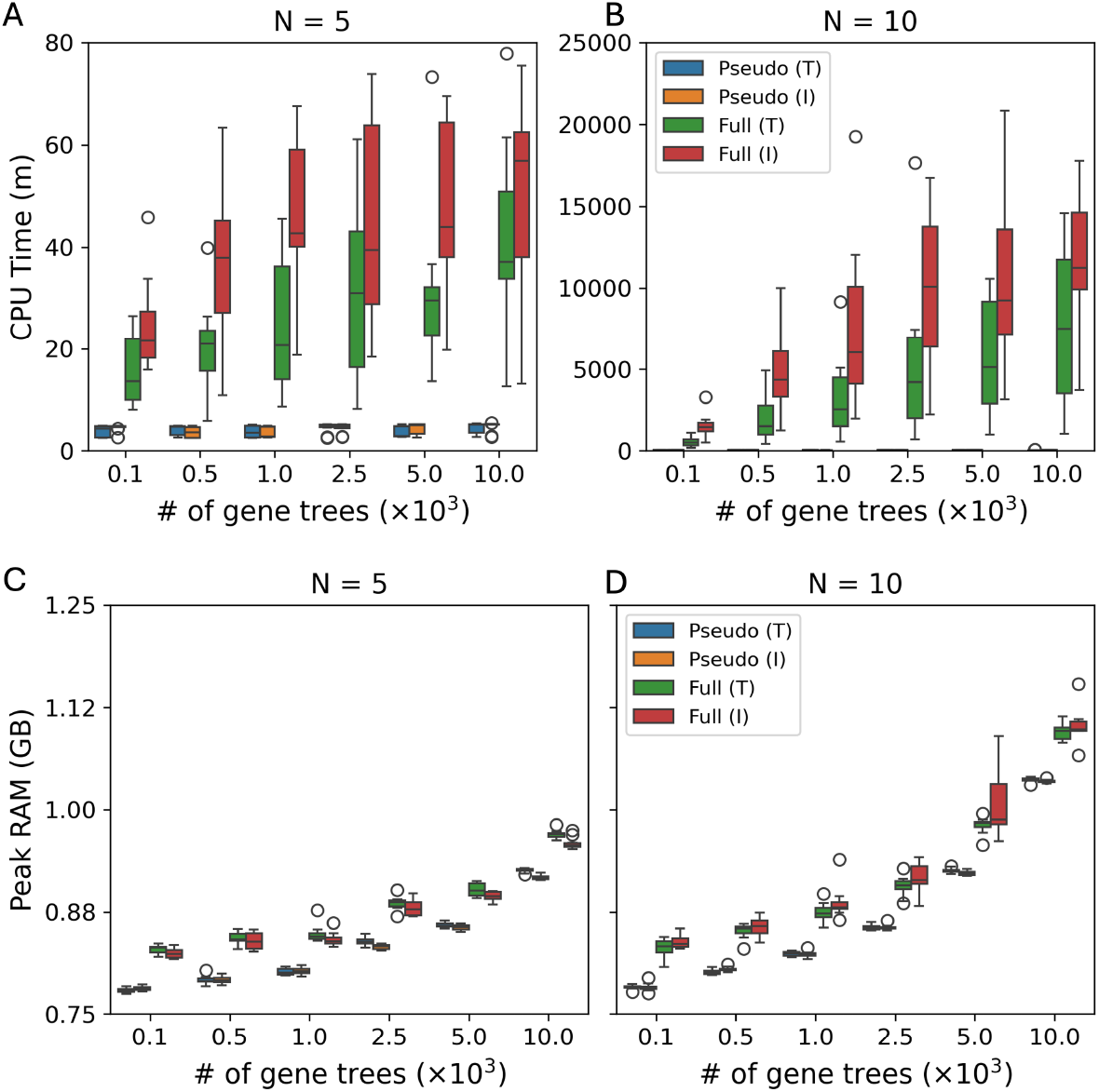
(A, B) CPU time required (in minutes) to complete 1,000,000 steps of the MCMC simulation on ground-truth gene trees (T) and inferred gene trees (I). (C, D) Peak RAM usage (in GB) during the MCMC simulation.

**Figure 13:**
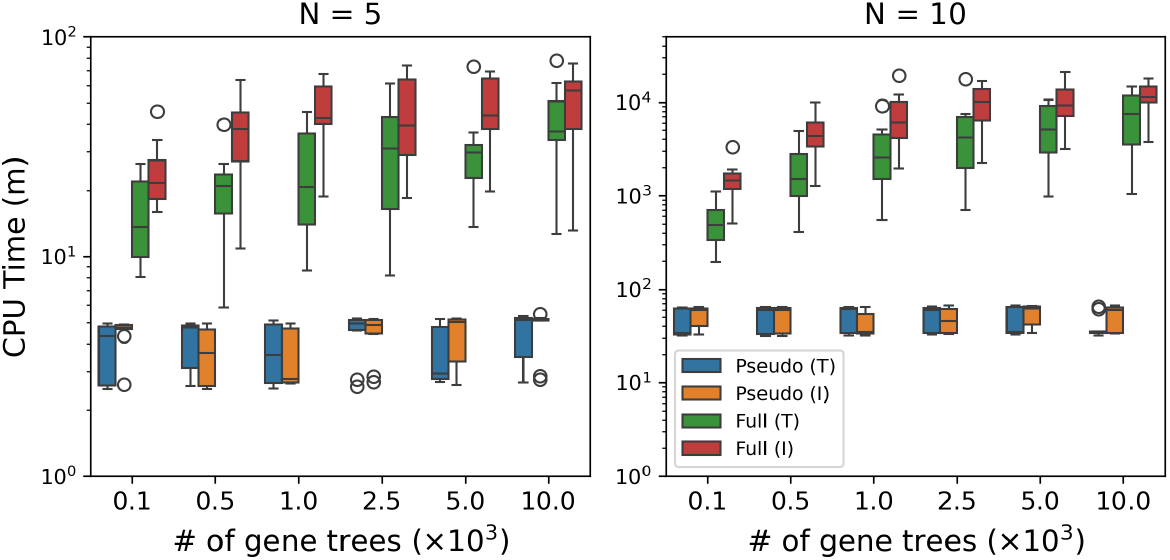
Logarithm of CPU time required (in minutes) to complete 1,000,000 steps of the MCMC simulation ground truth gene trees (T), and inferred gene trees (I) as a function of the number of gene trees (x-axis) and number of species (*N*).

The peak RAM usage of both methods is similar with full-likelihood-based analyses, requiring approximately 0.1 GB more RAM than pseudo-likelihood-based ones (Fig. 12C, D). All comparisons shown have been performed by running inference with a single thread on an Intel(R) Xeon(R) Gold 5220R CPU @ 2.20GHz.

### 3.3 Biological data analysis

We analyzed two biological phylogenomic datasets in this study. The first dataset comprised 233 alignments of ultra-conserved genetic elements (UCEs) from 32 taxa from order *Testudines* and 6 outgroup species (*H. sapiens, S. punctatus, A. carolinensis, P. molurus, C. porosus*, and *G. gallus*) [5]; we refer to this dataset as the **Turtles** dataset. The second dataset comprised 581 alignments from 83 taxa in the genus *Aphonopelma*; we refer to this dataset as the **Tarantulas** dataset.

For the **Turtles** dataset, all 900 sampled topologies were identical (Fig. 14) and had Robinson-Foulds distance of two with respect to the topology inferred by MrBayes [30] (from the concatenation of the 233 alignments) from the original study [5]. The discrepancy is caused by the placement of the *Sphenodon* outgroup. The branch length estimates obtained from our analyses deviated from those inferred in the original study, with only 22 out of 72 branches that had support in our sampled topologies having the branch length inferred in the original study within the 95% Bayesian credible interval. It is important to note here that the original analysis of the authors using MrBayes does not account for ILS explicitly, as MrBayes does not employ the multispecies coalescent model, whereas the analyses using our method do.

**Figure 14:**
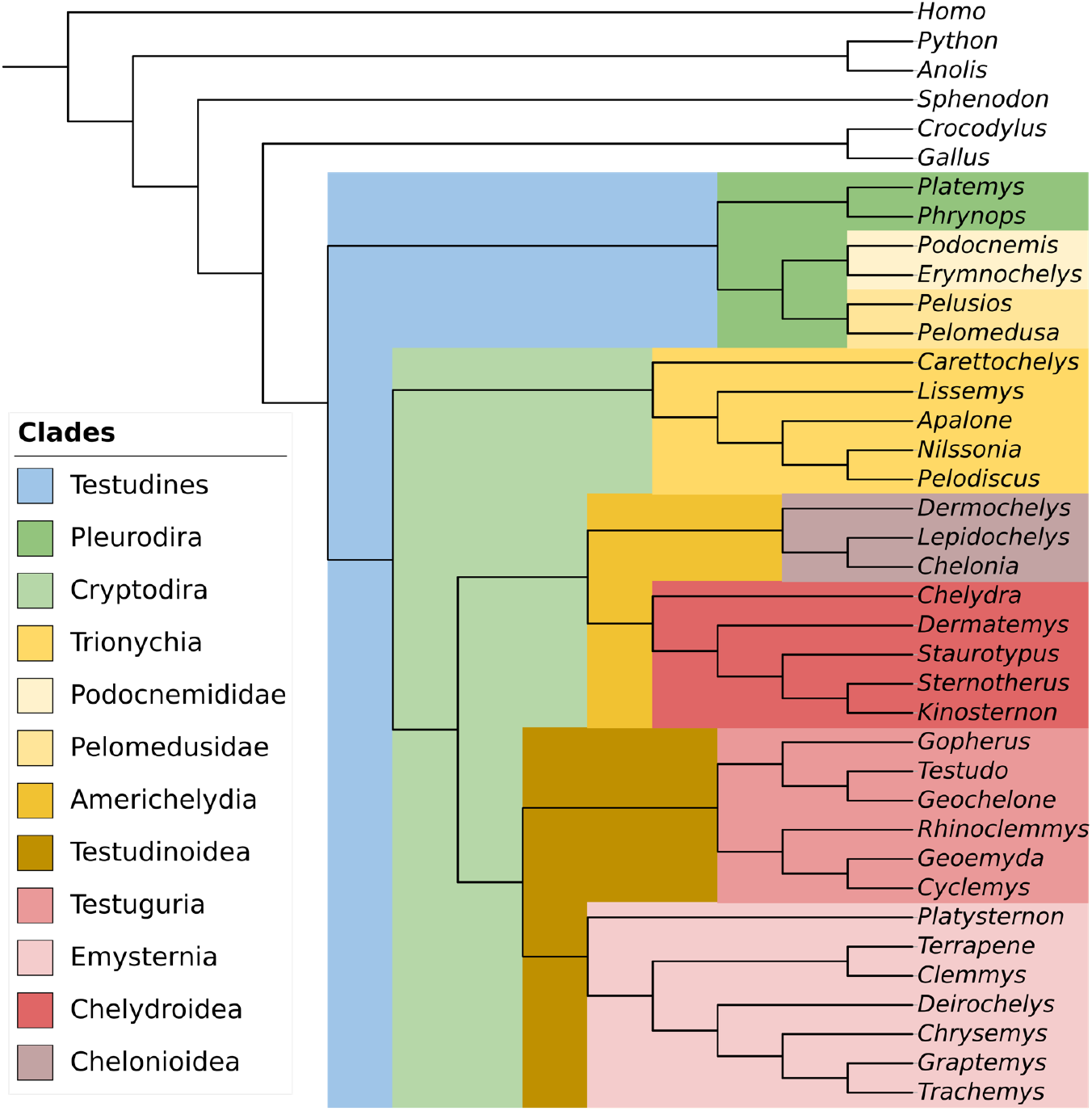
A cladogram of the maximum a posteriori (MAP) estimate of the species tree for the **Turtles** dataset inferred from 233 UCE trees. Major clades in order *Testudines* are highlighted with colors.

Analysis of this dataset has required a total of 70.83 CPU hours utilizing a single thread on an AMD EPYC 7642 48-Core Processor @ 2.30 GHz.

For the **Tarantulas** dataset (Fig. 15), of the 2,235 sampled topologies, 1,199 had Robinson-Foulds distance of 14 and 1,036 had Robinson-Foulds distance of 16 to the ASTRAL tree inferred in the original study [7]. The topology for the paraphyletic species group *Aphonopelma iodius* matches the one previously reported [7]. Discrepancies between the inferred trees and the ones reported in the original study arise due to the placement of the outgroup species, resolution of polytomies present in the ASTRAL tree, and topologies for the individual species (collapsed in Fig. 15). Of the 154 branches in the ASTRAL inferred species tree that had support in the sampled topologies, 70 had their branch length values estimated by ASTRAL within the 95% credible intervals.

**Figure 15:**
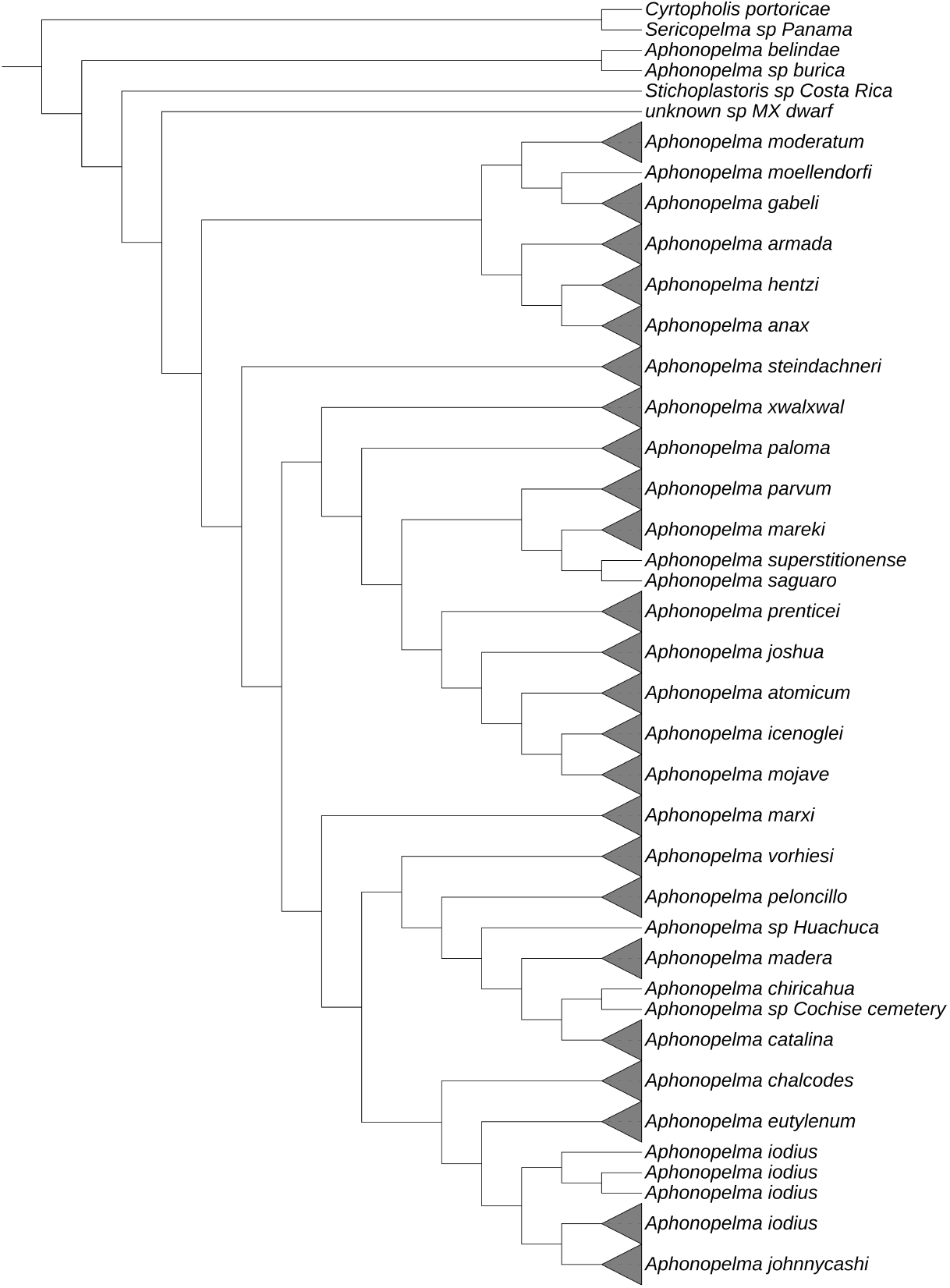
A cladogram of the maximum a posteriori (MAP) estimate of the species tree for the **Tarantulas** dataset inferred from 581 gene trees.

## 4 Discussion

Our theoretical and empirical analyses demonstrate the pseudo-likelihood provides a viable alternative to full likelihood for Bayesian phylogenomic inference tasks. It shows strong convergence guarantees with respect to the species tree topology supported by empirical evaluation in both idealized scenario where ground-truth gene trees are available and in a more realistic scenario where gene trees are estimated from the sequence data, although the impact of gene tree estimation error was only validated for low error rates (*<* 5%). Additionally, posterior distributions inferred based on pseudo-likelihood match those inferred based on full likelihood after appropriate centering. Furthermore, our observations on the required compute time indicate that the use of pseudo-likelihood can provide up to 100x speedup when compared with full-likelihood-based calculations. This is noteworthy given that the scale of biological datasets [5, 7] can completely prohibit Bayesian analyses under full likelihood. Additionally, since pseudo-likelihood-based inference speeds up individual proposal evaluations in the MCMC sampling procedure, it can in theory be paired with an improved proposal scheme [8, 41] to provide additional improvements to the computational cost.

However, as indicated by the theoretical analysis, branch length credible intervals do not match confidence intervals at the nominal level. While in theory the mismatch can result in both lower and higher empirical coverage, we note that in our experiments the coverage was always lower than the nominal value, that is, pseudo-likelihood-based Bayesian inference tends to be overconfident. Results on the biological data provide further evidence that the credible intervals estimated from pseudo-likelihood-based MCMC can be overconfident. These observations motivate a follow up investigation on whether there is a way to bound or correct this discrepancy, in particular by increasing the size of the sample derived from the MCMC run.

Finally, while species tree inference remains an important task, certain biological processes such as hybridization [1] and horizontal gene transfer can give rise to evolutionary histories that require inference of phylogenetic networks. The multispecies network coalescent (MSNC) has been previously proposed as a framework for statistical inference of phylogenetic species networks [39, 36]. However, computing full likelihood under the MSNC model is significantly more expensive when compared to species tree inference under the MSC model [42]. Thus, maximum pseudo-likelihood approaches have been proposed for the network inference problem under the MSNC [40, 31, 43]. However, suitability of pseudo-likelihood for Bayesian inference under the MSNC has not been investigated. Therefore, given the potential impact on the inference speed and promising results for species tree inference, we aim to expand current work to the network inference problem.

## 5 Acknowledgments

This work was in part supported by the NSF grants DMS/NIGMS-2153704 and DBI-2030604.

## References

[1] Nicholas H Barton and Godfrey M Hewitt. Analysis of hybrid zones. Annual review of Ecology and Systematics, pages 113–148, 1985.

[2] Natalia Bochkina. Bernstein–von Mises theorem and misspecified models: A review. Foundations of Modern Statistics, pages 355–380, 2019.

[3] David Bryant, Remco Bouckaert, Joseph Felsenstein, Noah A Rosenberg, and Arindam Roy-Choudhury. Inferring species trees directly from biallelic genetic markers: bypassing gene trees in a full coalescent analysis. Molecular Biology and Evolution, 29(8): 1917–1932, 2012.

[4] Samantha R Cook, Andrew Gelman, and Donald B Rubin. Validation of software for Bayesian models using posterior quantiles. Journal of Computational and Graphical Statistics, 15(3): 675–692, 2006.

[5] Nicholas G Crawford, James F Parham, Anna B Sellas, Brant C Faircloth, Travis C Glenn, Theodore J Papenfuss, James B Henderson, Madison H Hansen, and W Brian Simison. A phylogenomic analysis of turtles. Molecular Phylogenetics and Evolution, 83: 250–257, 2015.

[6] James H Degnan and Noah A Rosenberg. Discordance of species trees with their most likely gene trees. PLoS Genetics, 2(5):e68, 2006.

[7] Chris A Hamilton, Alan R Lemmon, Emily Moriarty Lemmon, and Jason E Bond. Expanding anchored hybrid enrichment to resolve both deep and shallow relationships within the spider tree of life. BMC Evolutionary Biology, 16: 1–20, 2016.

[8] Sebastian Hohna and Alexei J Drummond. Guided tree topology proposals for Bayesian phylogenetic inference. Systematic Biology, 61(1): 1–11, 2012.

[9] Richard R Hudson. Generating samples under a Wright–Fisher neutral model of genetic variation. Bioinformatics, 18(2): 337–338, 2002.

[10] T. Jukes and C. Cantor. Evolution of protein molecules. In H.N. Munro, editor, Mammalian Protein Metabolism, pages 21–132. Academic Press, NY, 1969.

[11] Joshua A Justison, Claudia Solis-Lemus, and Tracy A Heath. SiPhyNetwork: An R package for simulating phylogenetic networks. Methods in Ecology and Evolution, 14(7): 1687–1698, 2023.

[12] Subha Kalyaanamoorthy, Bui Quang Minh, Thomas KF Wong, Arndt Von Haeseler, and Lars S Jermiin. ModelFinder: fast model selection for accurate phylogenetic estimates. Nature Methods, 14(6): 587–589, 2017.

[13] BJK Kleijn and AW van der Vaart. The Bernstein-Von-Mises theorem under misspecification. Electronic Journal of Statistics, 6: 354–381, 2012.

[14] Laura S Kubatko, Bryan C Carstens, and L Lacey Knowles. STEM: species tree estimation using maximum likelihood for gene trees under coalescence. Bioinformatics, 25(7): 971–973, 2009.

[15] Meng Li, Zejian Liu, Cheng-Han Yu, and Marina Vannucci. Semiparametric Bayesian Inference for Local Extrema of Functions in the Presence of Noise. Journal of the American Statistical Association, 119(548): 3127–3140, 2024.

[16] Liang Liu, Lili Yu, and Scott V Edwards. A maximum pseudo-likelihood approach for estimating species trees under the coalescent model. BMC Evolutionary Biology, 10: 1–18, 2010.

[17] Wayne P Maddison. Gene trees in species trees. Systematic Biology, 46(3): 523–536, 1997.

[18] Siavash Mirarab, Luay Nakhleh, and Tandy Warnow. Multispecies coalescent: theory and applications in phylogenetics. Annual Review of Ecology, Evolution, and Systematics, 52(1):247–268, 2021.

[19] Siavash Mirarab, Rezwana Reaz, Md S Bayzid, Théo Zimmermann, M Shel Swenson, and Tandy Warnow. Astral: genome-scale coalescent-based species tree estimation. Bioinformatics, 30(17):i541–i548, 2014.

[20] Matthew Andres Moreno, Jeet Sukumaran, and Mark T Holder. Dendropy 5: a mature python library for phylogenetic computing. arXiv preprint arXiv:2405.14120, 2024.

[21] Luay Nakhleh. Computational approaches to species phylogeny inference and gene tree reconciliation. Trends in Ecology & Evolution, 28(12): 719–728, 2013.

[22] Lam-Tung Nguyen, Heiko A Schmidt, Arndt Von Haeseler, and Bui Quang Minh. IQ-TREE: a fast and effective stochastic algorithm for estimating maximum-likelihood phylogenies. Molecular Biology and Evolution, 32(1): 268–274, 2015.

[23] Huw A Ogilvie, Remco R Bouckaert, and Alexei J Drummond. StarBEAST2 brings faster species tree inference and accurate estimates of substitution rates. Molecular Biology and Evolution, 34(8): 2101–2114, 2017.

[24] Huw A Ogilvie, Joseph Heled, Dong Xie, and Alexei J Drummond. Computational performance and statistical accuracy of *BEAST and comparisons with other methods. Systematic Biology, 65(3): 381–396, 2016.

[25] Jingwen Pei and Yufeng Wu. STELLS2: fast and accurate coalescent-based maximum likelihood inference of species trees from gene tree topologies. Bioinformatics, 33(12): 1789–1797, 2017.

[26] Andrew Rambaut and Nicholas C Grass. Seq-Gen: an application for the Monte Carlo simulation of DNA sequence evolution along phylogenetic trees. Bioinformatics, 13(3): 235–238, 1997.

[27] Bruce Rannala and Ziheng Yang. Bayes estimation of species divergence times and ancestral population sizes using dna sequences from multiple loci. Genetics, 164(4): 1645–1656, 2003.

[28] David F Robinson and Leslie R Foulds. Comparison of phylogenetic trees. Mathematical biosciences, 53(1–2): 131–147, 1981.

[29] David F Robinson and Leslie R Foulds. Comparison of weighted labelled trees. In Combinatorial Mathematics VI: Proceedings of the Sixth Australian Conference on Combinatorial Mathematics, Armidale, Australia, August 1978, pages 119–126. Springer, 2006.

[30] Fredrik Ronquist, Maxim Teslenko, Paul Van Der Mark, Daniel L Ayres, Aaron Darling, Sebastian Hohna, Bret Larget, Liang Liu, Marc A Suchard, and John P Huelsenbeck. MrBayes 3.2: efficient Bayesian phylogenetic inference and model choice across a large model space. Systematic Biology, 61(3): 539–542, 2012.

[31] Claudia Solís-Lemus and Cécile Ané. Inferring phylogenetic networks with maximum pseudo-likelihood under incomplete lineage sorting. PLoS Genetics, 12(3):e1005896, 2016.

[32] Cuong Than, Derek Ruths, and Luay Nakhleh. PhyloNet: a software package for analyzing and reconstructing reticulate evolutionary relationships. BMC Bioinformatics, 9: 1–16, 2008.

[33] Aad van der Vaart. Asymptotic Statistics. Cambridge University Press, 1998.

[34] Yaxuan Wang, Huw A Ogilvie, and Luay Nakhleh. Practical speedup of Bayesian inference of species phylogenies by restricting the space of gene trees. Molecular Biology and Evolution, 37(6): 1809–1818, 2020.

[35] Yixin Wang and David M. Blei. Frequentist Consistency of Variational Bayes. Journal of the American Statistical Association, 114(527):1147–1161, jul 2019.

[36] Dingqiao Wen, Yun Yu, and Luay Nakhleh. Bayesian inference of reticulate phylogenies under the multispecies network coalescent. PLoS Genetics, 12(5):e1006006, 2016.

[37] Dingqiao Wen, Yun Yu, Jiafan Zhu, and Luay Nakhleh. Inferring phylogenetic networks using PhyloNet. Systematic Biology, 67(4): 735–740, 2018.

[38] Yufeng Wu. Coalescent-based species tree inference from gene tree topologies under incomplete lineage sorting by maximum likelihood. Evolution, 66(3): 763–775, 2012.

[39] Yun Yu, Jianrong Dong, Kevin J Liu, and Luay Nakhleh. Maximum likelihood inference of reticulate evolutionary histories. Proceedings of the National Academy of Sciences, 111(46): 16448–16453, 2014.

[40] Yun Yu and Luay Nakhleh. A maximum pseudo-likelihood approach for phylogenetic networks. BMC Genomics, 16: 1–10, 2015.

[41] Chi Zhang, John P Huelsenbeck, and Fredrik Ronquist. Using parsimony-guided tree proposals to accelerate convergence in Bayesian phylogenetic inference. Systematic Biology, 69(5):1016–1032, 2020.

[42] Jiafan Zhu, Xinhao Liu, Huw A Ogilvie, and Luay K Nakhleh. A divide-and-conquer method for scalable phylogenetic network inference from multilocus data. Bioinformatics, 35(14):i370–i378, 2019.

[43] Jiafan Zhu and Luay Nakhleh. Inference of species phylogenies from bi-allelic markers using pseudo-likelihood. Bioinformatics, 34(13):i376–i385, 2018.

